# Optogenetic Regulation of EphA1 RTK Activation and Signaling

**DOI:** 10.1101/2024.02.06.579139

**Authors:** Anna I Wurz, Kevin Zheng, Robert M. Hughes

## Abstract

Eph receptors are ubiquitous class of transmembrane receptors that mediate cell-cell communication, proliferation, differentiation, and migration. EphA1 receptors specifically play an important role in angiogenesis, fetal development, and cancer progression; however, studies of this receptor can be challenging as its ligand, ephrinA1, binds and activates several EphA receptors simultaneously. Optogenetic strategies could be applied to circumvent this requirement for ligand activation and enable selective activation of the EphA1 subtype. In this work, we designed and tested several iterations of an optogenetic EphA1 - Cryptochrome 2 (Cry2) fusion, investigating their capacity to mimic EphA1-dependent signaling in response to light activation. We then characterized the key cell signaling target of MAPK phosphorylation activated in response to light stimulation. The optogenetic regulation of Eph receptor RTK signaling without the need for external stimulus promises to be an effective means of controlling individual Eph receptor-mediated activities and creates a path forward for the identification of new Eph-dependent functions.

## Introduction

Erythropoietin-producing hepatocellular carcinoma (Eph) receptors are the largest family of receptor tyrosine kinases (RTK) with 14 different types of this transmembrane protein (Arvanitis & Davy, 2008). Each Eph receptor includes an extracellular domain, comprising of the ligand-binding domain on the N-terminal, a cysteine-rich region, and two consecutive fibronectin type-III domains that are adjacent to the transmembrane region of the Eph receptor (**Fig. 1A**) (Hughes & Virag, 2020) and an intracellular portion consisting of a juxtamembrane region, followed by a kinase domain, a sterile alpha motif (SAM) domain, and ending in a PDZ-binding motif at the C-terminus. Each of the two classes of Eph receptors, EphA (A1-A8, A10) and EphB (B1-5), is classified by the binding of their corresponding ligands, ephrinA and ephrinB, respectively (Arvanitis & Davy, 2008; Liang et al., 2019). EphrinA and B both contain an extracellular receptor-binding domain that interacts with the ligand-binding domain on the Eph receptor; however, ephrinA is anchored to the membrane by a glycosylphosphatidylinositol (GPI) post-translational modification, and ephrinB has a transmembrane domain with an intracellular PDZ-binding motif on the C-terminus (Darling & Lamb, 2019; Liang et al., 2019).

**Figure 1.**
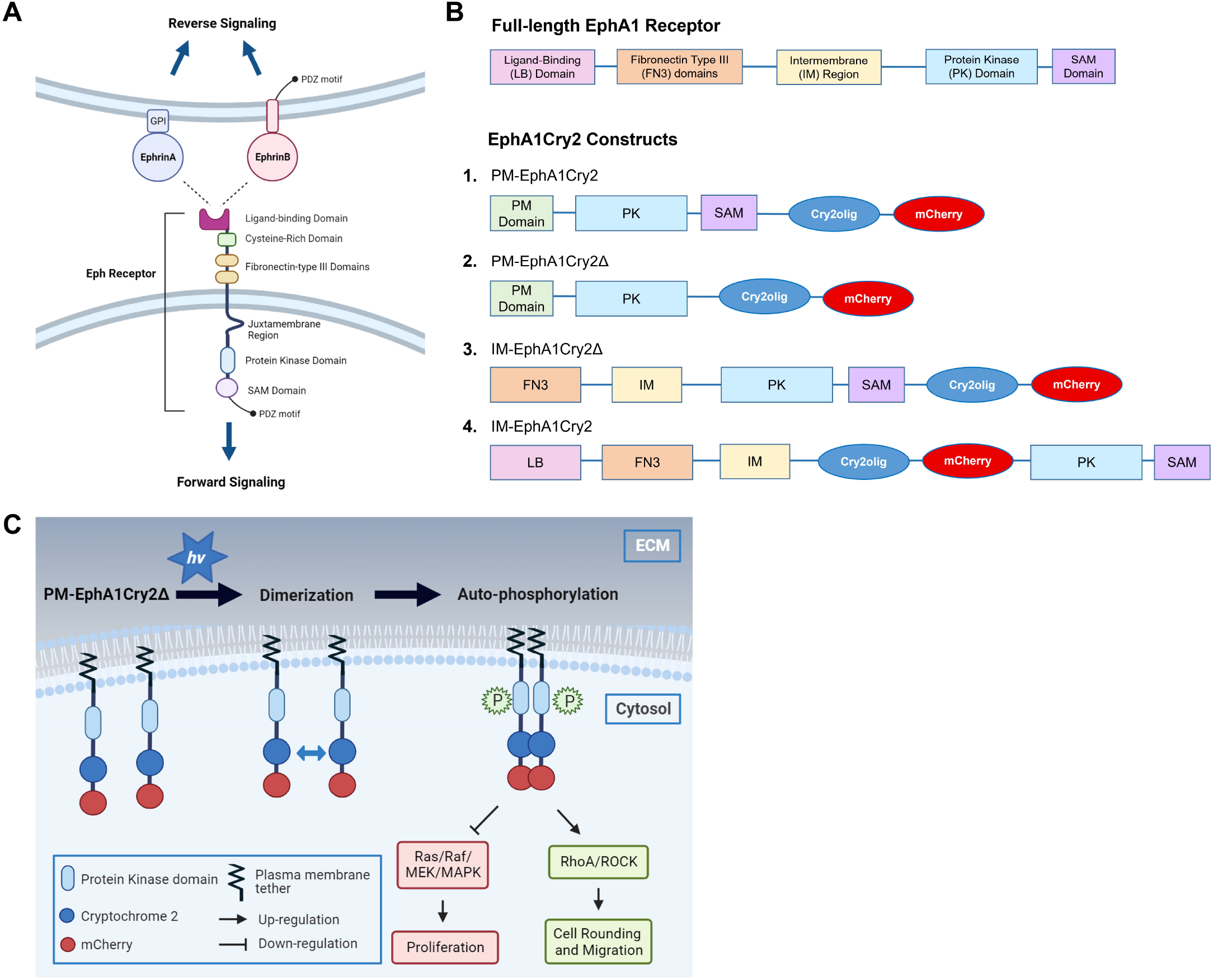
**(A)** Schematic of Eph receptor and ephrin and their bi-directional signaling. Ephrin binds to the extracellular ligand-binding domain of the Eph receptor expressed on a neighboring cell. This interaction activates reverse signaling by the ephrin ligand in one cell and forward signaling by the Eph receptor in the other cell. **(B)** Schematic of full-length wild-type EphA1 receptor domains (top) and EphA1Cry2 constructs (bottom, labeled 1-4). **(C)** Schematic of the hypothesis of PM-EphA1Cry2 construct activation and signaling. When exposed to blue light, the constructs oligomerize via Cry2 interactions and the adjacent protein kinase domains auto-phosphorylate each other, activating downstream phosphorylation of signaling proteins that induce cell rounding and migration through the RhoA/ROCK pathway and inhibit proliferation through the Ras/Raf/MEK/MAPK pathway.

The ephrin/Eph system incorporates bi-directional signaling pathway, triggered by the binding of ephrin to the Eph’s ligand-binding domain (**Fig. 1A**). Forward signaling activates kinase signaling through autophosphorylation of the tyrosine kinase domain, leading to downstream signaling. Reverse signaling is activated through cell surface displayed ephrin ligands and controls ephrin-specific pathways in the host cell. This distinctive type of signaling has implications for cell-cell communication, including cell repulsion, migration, differentiation, and proliferation (Barquilla & Pasquale, 2015; Darling & Lamb, 2019; Gucciardo et al., 2014; Hughes & Virag, 2020; Lai & Ip, 2009; Liang et al., 2019; Lisabeth et al., 2013). In neural development for example, forward signaling mediates axonal growth cone collapse, and reverse signaling induces growth cone survival, which in tandem can regulate migration of axons to their target destinations (Lai & Ip, 2009; Lisabeth et al., 2013).

These complex, overlapping signals make it difficult to determine the effects of individual Eph receptor activation and disentangle subsequent downstream signaling. As a result, Eph and ephrin signaling crosstalk is not completely understood, and the reports of the impacts of Eph/ephrin signaling activation are sometimes contradictory (Ieguchi & Maru, 2019). To simplify the complex web of signaling pathways, synthetic biology strategies for the control of specific Eph receptor types could more precisely delineate Eph signaling outcomes, leading to a better understanding of the impacts of Eph-dependent cell signaling cascades (Locke et al., 2017). Optogenetics comprises one such strategy that could be applied to the control of Eph receptor signaling.

In this work, we use cryptochrome 2 (Cry2), a blue light-dependent photoreceptor found in *Arabidopsis thaliana* (Kennedy et al., 2010) to create a light responsive EphA1 receptor mimic. Cry2 has been used in numerous protein fusion experiments due to its ability to homo-oligomerize in the cell cytosol in response to blue light stimulation (Bugaj et al., 2015; Hernández-Candia et al., 2021) and enables direct regulation of various cellular processes solely by control with light (Duan et al., 2017; Hughes, 2018; Kennedy et al., 2010). In prior studies of optogenetic control of Eph receptor domains, Locke et. al. described an optogenetic EphB2 receptor, named OptoEphB2, consisting of only the intracellular domains of EphB2 fused C-terminal to Cry2 and the fluorescent protein, mCherry (Locke et al., 2017). In these studies, a mutant of Cry2, Cry2olig, was used to improve light-initialized oligomerization (Taslimi et al., 2014). For this study, EphA1 was selected as the receptor of interest due to its role in cell adhesion and migration, cancer progression, and Alzheimer’s disease (Carrasquillo et al., 2011; Gucciardo et al., 2014; Ieguchi & Maru, 2019; Naj et al., 2011; Yamazaki et al., 2009). Although EphA1 was the first Eph receptor discovered in 1987 (Hirai et al., 1987), its specific function in different cell types has yet to be fully investigated (Ieguchi & Maru, 2019). Its complementary ligand, ephrinA1, has also been studied for its reverse signaling effect, which similarly promotes cell de-adhesion and migration (Yang et al., 2018). In previous studies of wild-type EphA1, ephrinA1-Fc stimulation was required to activate the receptor and produce cellular response (Yamazaki et al., 2009). To create a light-activated EphA1 that functions in the absence of ephrinA1 ligand stimulation, we coupled Cry2olig with various truncations EphA1 and investigated their ability to oligomerize and undergo RTK-dependent autophosphorylation.

## Material and Methods

### EphA1Cry2 construct design

Forward and reverse primers for the four EphA1 constructs were designed and PCR amplified using human EphA1 template (P21709). Construct PM-EphA1Cry2 and PM-EphA1Cry2Δ primers included a plasma membrane tether at the C-terminus (PMa amino acid sequence: MGCIKSKRKDNLNDDE (Resh, 1999; Thaa et al., 2015); PMb amino acid sequence: MGSSKSKPK (Locke et al., 2017). Select regions of EphA1 (excluding the extracellular regions of IM-EphA1Cry2 construct) were cloned into Cry2olig-mCherry (Addgene: #60032) using NheI and BmtI restriction sites. The extracellular region of IM-EphA1Cry2 was cloned into Cry2olig-mCherry using BsrgI and NotI. The constructs were transformed into competent *E. Coli*, purified, and characterized via Sanger sequencing.

### Cell culture preparation

HEK293T cells and Neuro2a cells were cultured in Dulbecco’s Modified Eagle Medium (DMEM) (Gibco) supplemented with 10% fetal bovine serum (FBS) and 1% Penicillin/Streptomycin (PenStrep) (Gibco) in an incubator at 37°C and 5% CO_2_. Cells were plated on glass-bottom dishes for imaging or 6-well plates for lysis and transfected with each construct using Calfectin (Signa Gen Laboratories). For ERK and Akt immunostaining, HEK293T cells were serum-starved in DMEM with 0.1% FBS for 2 hours prior to light exposure and lysis.

### Neuro2a differentiation

Neuro2a cells were seeded at 60% confluency in 1 mL DMEM with 10% FBS and 1% PenStrep on 35 mm glass-bottom dishes. The media was replaced after 24 hours with DMEM with 1% FBS and 20 µM retinoic acid. The media was replaced daily for 3 days. On the third day, cells were transfected with either PMb-EphA1Cry2 or Cry2olig using Calfectin (Signa Gen Laboratories) in differentiation media and used for imaging the next day. Cells were serum-starved in 1x PBS for 2-4 hours prior to imaging.

### Kinase translocation reporter subcloning and experiment

ERK-KTR-mNeonGreen was subcloned from Addgene plasmid #129631 (Chavez-Abiega, 2022) into a phCMV vector (Genelantis). HEK293T cells were transfected with 1000 ng of ERK-KTR or co-transfected with 1000 ng of each construct using Calfectin (Signa Gen Laboratories). The following day, cells were serum-starved with Leibovitz’s L-15 medium for 1-2 hours. Cells were then incubated in 1 mL of 20 mM Hoechst 33342 nuclear stain in DPBS, washed with 1 mL DPBS, and placed back in L15 media for imaging. For ERK-KTR only cells, a 10-minute pre-stimulus period was imaged, then cells were stimulated with 500 ng/mL of epidermal growth factor (EGF).

### Microscopy and image analysis

HEK293T cells and Neuro2a cells were imaged on a Leica DMi8 Live Cell Imaging System, equipped with an OKOLab stage-top live cell incubation system, LASX software, Leica HCX PL APO 63×/1.40 to 0.60 numerical objective oil objective, Lumencor LED light engine, CTRadvanced+ power supply, and a Leica DFC900 GT camera. The mCherry channel (553 nm, 50% power, 200 ms exposure) was used for pre, during, and post-light visualization. GFP channel (480 nm, 50% power, 50 ms exposure) was used for Cry2 activation. Images were captured in 30 sec intervals.

Live-cell confocal images of HEK293T cells were collected on Zeiss LSM 700 laser scanning microscope equipped with a stage-top incubator and ZEN Black 2012 software. Images were captured on the mCherry channel (553 nm) and GFP channel (480 nm) for blue light illumination in 30 sec intervals. For ERK-KTR cell experiments, ERK-KTR was visualized on the GFP channel (5% power), and the nuclear stain was visualized on the DAPI channel (395 nm, 5% power). All images were analyzed using Fiji Image J software.

### Western Blotting

Transfected HEK293T cells were lysed in M-PER (Thermo Scientific) with 1X HALT protease & phosphatase inhibitor (Thermo Scientific) either in the dark or immediately after illumination with blue light using a homemade LED lightbox for the specified time periods. After 10 min of shaking, total cell lysates were centrifuged for 15 min at 4°C. Laemmli SDS sample buffer (Alfa Aesar) was added to supernatants and incubated at 65 °C for 10 min. Cell lysates were separated by 10% SDS-PAGE gel electrophoresis and transferred to a polyvinylidene fluoride membrane (70V for 90 min on ice). Membranes were blocked with 5% bovine serum albumin (BSA) in 1x Tris-buffered saline (TBS) with 1% Tween (TBST) for 1 hour, room temperature. Blots were then incubated with primary antibody in a 1:1000 dilution with 5% BSA in TBST overnight at 4°C: tyrosine phosphorylation using phosphotyrosine mouse monoclonal Ig pY99 (Santa Cruz Biotechnologies); mCherry expression using anti-mCherry rabbit polyclonal Ig (Rockland Inc.); GAPDH expression using GAPDH mouse monoclonal Ig (Invitrogen); MAPK phosphorylation using p-p44/42 MAPK T202/Y204 rabbit Ig (Cell Signaling Technologies); total ERK using p44/42 MAPK (ERK1/2) rabbit Ig (Cell Signaling Technologies). Blots were washed with TBST 3x for 5 min and incubated in corresponding secondary antibody for 1 hour room temperature: goat anti-rabbit antibody (Invitrogen) or rabbit anti-mouse antibody (Invitrogen). The membranes were again washed with TBST 3x for 5 min, incubated with a chemiluminescent substrate for 5 min, and imaged with an Azure cSeries imaging station.

### Design of EphA1Cry2 constructs

Four optogenetic EphA1 receptor constructs were created and screened for their membrane localization and their light-activated clustering and RTK tyrosine autophosphorylation properties (**Fig. 1B**). We based the first optogenetic construct on previous optogenetic receptor designs (**Fig. 1B.1**) (Locke et al., 2017). This construct (**PM-EphA1Cry2**) contains a plasma membrane localization sequence (labeled PM) fused to the N-terminus of the intracellular protein kinase domain and SAM domain of the EphA1 receptor; Cry2olig is tethered to the C-terminus via a glycine-rich linker sequence, followed by mCherry fluorescent protein. Two separate PM localization sequences were used (denoted PMa and PMb): PMa represents a Lyn myristoylation domain with the amino acid sequence MGCIKSKRKDNLNDDE and PMb represents the membrance localization sequence MGSSKSKPK (Locke et al., 2017). The second construct (**PM-EphA1Cry2**Δ) is identical to the PM-EphA1Cry2 construct except the SAM domain was removed; Δ indicates the absence of the SAM domain (**Fig 1B.2**). The third and fourth constructs (**IM-EphA1Cry2**Δ and **IM-EphA1Cry2**) include the intermembrane (IM) region (**Fig 1B.3 and 4**). **IM-EphA1Cry2**Δ contains the intracellular protein kinase and SAM domains with the additional extracellular fibronectin type III domain (**Fig. 1B.3**) without the external ligand binding domain (indicated by Δ in the name). **IM-EphA1Cry2** contains all intracellular and extracellular receptor domains, with Cry2olig-mCherry was fused between the intermembrane region and the protein kinase domain (**Fig. 1B.4**). The proposed scheme for optogenetic EphA1 receptor activation and signaling is detailed in **Figure 1C**: upon blue light exposure, Cry2olig-mediated oligomerization should induce RTK clustering and autophosphorylation and subsequent activation of downstream signaling.

Optogenetic EphA1 constructs were initially tested via transfection in HEK293T and subsequent activation and imaging on a widefield microscope (**Fig. 2A**). For each construct, the resulting images were analyzed to determine the extent of membrane localization and light-activated clustering in response to light. As anticipated, the cytosolic Cry2olig control showed high-order clustering in the cytosol after 5 min of blue light exposure compared to the cell pre-stimulus at 0 min (**Fig. 2A, row 1**). By contrast, inspection of plasma membrane localization revealed that the PMa-EphA1Cry2 construct was not well localized to the plasma membrane (**Fig. 2A, row 2**), whereas PMa-EphA1Cry2Δ was membrane localized (**Fig. 2D, row 3**). To quantify membrane localization and cluster formation at the membrane upon light stimulation, the fluorescent intensity was measured at the membrane and cytosol over time of light exposure (**Fig. 2C**). Both the PMa-EphA1Cry2 and PMa-EphA1Cry2Δ exhibited an increase in fluorescent intensity at the membrane and a decrease in fluorescent intensity in the cytosol (**Fig. 2D**). Conversely, the Cry2olig control exhibited an increase in fluorescent intensity in the cytosol and a decrease in intensity at the membrane, due to recruitment and localization of Cry2 oligomers within the cytosol (**Fig. 2D**). Construct clustering at the membrane did improve slightly for PMb-EphA1Cry2 (**Fig. 2B, row 1**), but PMb-EphA1Cry2Δ construct showed a notable improvement in localization and clustering upon light exposure (**Fig. 2, row 2; Fig. 2D**) compared to PMa-EphA1Cry2Δ. Both IM-EphA1Cry2 and IM-EphA1Cry2Δ construct exhibited membrane localization but did not show visible clustering in response to light activation (**Fig. 2B, row 3 and 4**). We also investigated the ability of our constructs to be activated with spatially restricted illumination and undergo dark reversion. Constructs PMa-EphA1Cry2, PMa-EphA1Cry2Δ, and PMb-EphA1Cry2Δ were confirmed to be spatially activated, and both PMa-EphA1Cry2Δ and PMb-EphA1Cry2Δ construct activation were reversible (**see Supporting Figure 1 and Supporting Movie 1**). In addition, we performed a cell rounding assay for constructs PMa-EphA1Cry2, PMa-EphA1Cry2Δ, and PMb-EphA1Cry2Δ as previous studies have found EphA1 receptor activation promotes cell rounding (Yamazaki et al., 2009). All three constructs exhibited a decrease in cell area after light stimulation compared to the Cry2olig control (**see Supporting Figure 2)**. Going forward, only constructs that showed a strong membrane-adjacent light activated clustering response were selected for further analysis.

**Figure 2.**
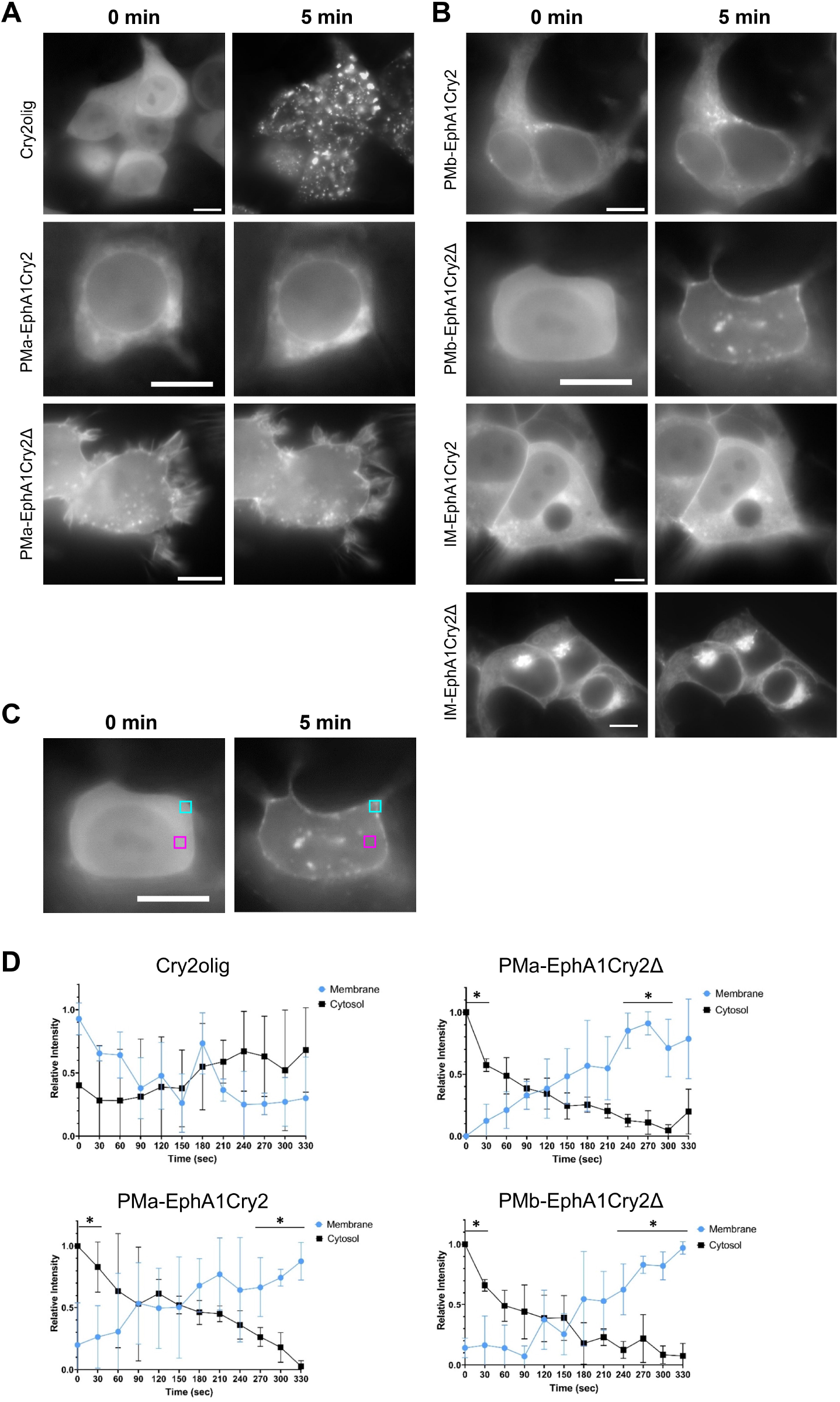
Optogenetic clustering of EphA1Cry2 constructs. HEK293T cells were transfected with **(A)** Cry2olig (top row), PMa-EphA1Cry2 (middle row), or PMa-EphA1Cry2Δ (bottom row); and **(B)** PMb-EphA1Cry2 (top row), PMb-EphA1Cry2Δ (second row), IM-EphA1Cry2 (third row), or IM-EphA1Cry2Δ (bottom row). **(A**,**B)** Widefield images of transfected cells were taken pre-illumination (0 min) and after blue light exposure (480 nm, 50 ms exposure, 30 sec intervals over 5 min). Images collected on mCherry channel (553 nm, 200 ms exposure). Scale bar: 10 µm. **(C)** Representative image of relative fluorescent intensity quantitation using regions of interest in the cytosol (magenta box, 2.5 × 2.5 μm) and at the membrane (cyan box, 2.5 × 2.5 μm) of a PMb-EphA1Cry2Δ transfected HEK293T cell. **(D)** Quantitation of the relative intensity of region of interests (C) in the cytosol (black) and at plasma membrane (blue) over 5 min (330 sec) of blue light exposure, taken every 30 sec. Data was collected for Cry2olig, PMa-EphA1Cry2, PMa-EphA1Cry2Δ, and PMb-EphA1Cry2Δ constructs using Fiji Image J and graphed using GraphPad Prism. Error bars represent standard deviation; n=3 (unpaired t-test, ^*^p < 0.05).

### Tyrosine phosphorylation in response to light activation

Next, we investigated global tyrosine phosphorylation to capture the extent of light-activated RTK activation of the selected constructs: PMa-EphA1Cry2, PMa-EphA1Cry2Δ, and PMb-EphA1Cry2Δ. Analysis of anti-phosphotyrosine (pTyr) western blots of light vs. Dark conditions for PMa-EphA1Cry2, PMa-EphA1Cry2Δ, and PMb-EphA1Cry2Δ constructs showed an upward trend in phosphorylation for each construct over time compared to no change in phosphorylation for Cry2olig (**Fig. 3A, B**). When performing an initial comparison blot between PMa- and PMb-EphA1Cry2, we noticed very light phosphorylation bands at the molecular weight of the PMb-EphA1Cry2 construct but a dark band at around 50 kDa that was not present in the PMa-EphA1Cry2 lanes (**Fig. 3C**). The lower molecular weight band is most likely due to degradation of the construct; as a result, PMb-EphA1Cry2 was not investigated further. For constructs PMa-EphA1Cry2, PMa-EphA1Cry2Δ, PMb-EphA1Cry2Δ, a slight decrease in tyrosine phosphorylation was observed between the 20 min and 30 min time points and smaller error for 30 min than the 20 min time point (**Fig. 3B**). Based on these data, we performed a separate western blot for tyrosine phosphorylation for cells in the dark or exposed to 30 min of blue light stimulation for the three constructs in Figure 3C (**Fig. 3D**). Each construct exhibited a significant log fold change increase of phosphorylation after light exposure compared to the Cry2olig control (**Fig. 3E**). Although PMb-EphA1Cry2Δ exhibited the lowest overall phosphotyrosine band intensity, it exhibited the lowest background phosphorylation with intensities at time 0 min comparable to Cry2olig intensities (**Fig. 3B**) and additionally showed the largest log fold change amongst the three EphA1Cry2 constructs (**Fig. 3E**). As a result, PMb-EphA1Cry2Δ was selected as the lead construct for further study.

**Figure 3.**
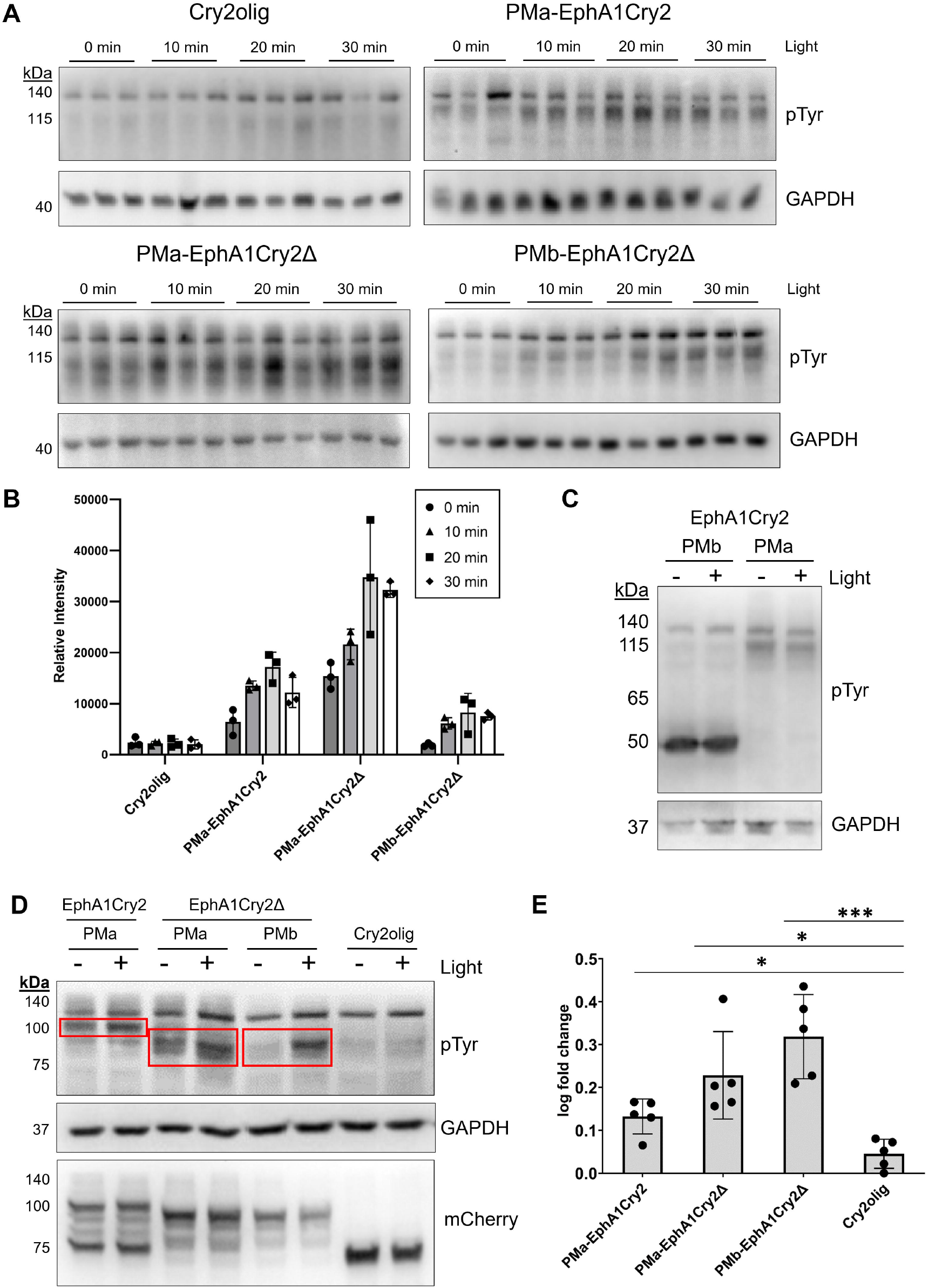
Analysis of EphA1Cry2 tyrosine phosphorylation. **(A)** HEK293T cells were transfected with either Cry2olig, PMa-EphA1Cry2, PMa-EphA1Cry2Δ, or PMb-EphA1Cry2Δ and lysed under dark conditions (0 min light) or after exposure to either 10, 20, or 30 min of LED blue light. Western blots of whole cell lysates were incubated with pTyr99 antibody for tyrosine phosphorylation and GAPDH antibody for total protein expression in triplicate. Bands at ∼130 kDa are endogenous bands. **(B)** Quantitation of normalized pTyr band intensities at the molecular weight of each construct (∼115 kDa) to GAPDH (A). Intensities were measured using Fiji ImageJ and plotted using GraphPad Prism. Error bars represent standard deviation; n=3. **(C)** HEK293T cells transfected with either PMa-EphA1Cry2 or PMb-EphA1Cry2 were lysed under dark conditions (-) or after 30 min of LED blue light exposure (+). Western blots of whole cell lysates were incubated with pTyr antibody for tyrosine phosphorylation and GAPDH antibody for total protein expression. **(D)** HEK293T cells transfected with either PMa-EphA1Cry2 or PMb-EphA1Cry2 were lysed under dark conditions (-) or after 30 min of LED blue light exposure (+). Western blots of whole cell lysates were incubated with pTyr antibody for tyrosine phosphorylation, GAPDH antibody for total protein expression, and mCherry for construct expression. Red boxes indicate bands at the construct molecular weight. **(E)** Quantitation of normalized pTyr band intensity to GAPDH (D), shown as the log fold change of light vs. dark condition. Error bars represent standard deviation, n=5 (unpaired t-test; ^*^p < 0.05, ^***^p < 0.001).

### EphA1Cry2 phosphorylates ERK upon light activation

EphA2 has been characterized as an inhibitor of proliferation by inhibition of the Ras/Raf/MEK/MAPK signaling cascade (Miao et al., 2000, 2001). Unlike other RTKs, EphA receptors function as the opposite of growth factors to inhibit the cascade that leads to an increase in proliferation (Miao et al., 2001). EphA2 has been proposed as a therapeutic target for cancer to due to this cascade, and upregulation could yield a decrease in the progression of certain cancers (Miao & Wang, 2012). Although EphA1 has also been hypothesized to promote tumorigenesis and metastasis (Ieguchi & Maru, 2019), EphA1 receptor activation of signaling proteins associated with cell proliferation is less well defined. To identify downstream signaling events specific to proliferation, we investigated whether light activation of EphA1Cry2 impacts phosphorylation of extracellular-signal regulated kinase (ERK), also called mitogen-activated protein kinases (MAPK), a known signaling target for EphA receptors (**Fig. 4**) (Miao et al., 2001). Interestingly, we found an increase in ERK phosphorylation upon PMb-EphA1Cry2Δ light activation over time with and without serum-starvation of cells via western blot (**Fig. 4A, B**). We also utilized an ERK kinase translocation reporter (KTR) to corroborate the western blot results (**Fig. 4C, D**). When ERK is phosphorylated by upstream receptor signaling proteins, p-ERK translocates to the nucleus and phosphorylates nuclear ERK-KTR. Phosphorylated ERK-KTR then translocates from the nucleus into the cytosol (**Fig. 4E**). The KTR translocation event can be quantified by measuring and then calculating the ratio of the fluorescent intensity within the cytosol to the nucleus (C/N ratio) of the associated mNeonGreen fusion. As a positive control, ERK-KTR was transfected in HEK293T cells and then treated with epidermal growth factor (EGF) to stimulate endogenous EGF receptors in the cell and promote phosphorylation of ERK. The cells transfected with ERK-KTR exhibited an immediate increase in C/N ratio after EGF stimulation, peaking at about 150 sec (**Fig. 4C, F**). This response is consistent with previously reported C/N changes of this reporter and other similar ERK reporters (Chavez-Abiega et al., 2022; Dessauges et al., 2022). By comparison, PMb-EphA1Cry2Δ light activation resulted in a steady increase in ERK activation over time of light exposure (**Fig. 4D, F; Supporting Movie 2**), which is consistent with the ERK phosphorylation western blot (**Fig. 4A, B**). Taken together, these results indicate that light-induced PMb-EphA1Cry2Δ construct activation stimulates downstream activation of ERK. Conversely, light activation of Cry2olig did not show a trend in C/N change over time (**Fig. 4D, F)**, also corroborating the western blot data (**Fig. 4A, B**). We note that the general trend of increasing ERK activity upon light simulation mirrors that of similar optogenetic RTK proteins, such as optoFGFR (Dessauges et al., 2022).

**Figure 4.**
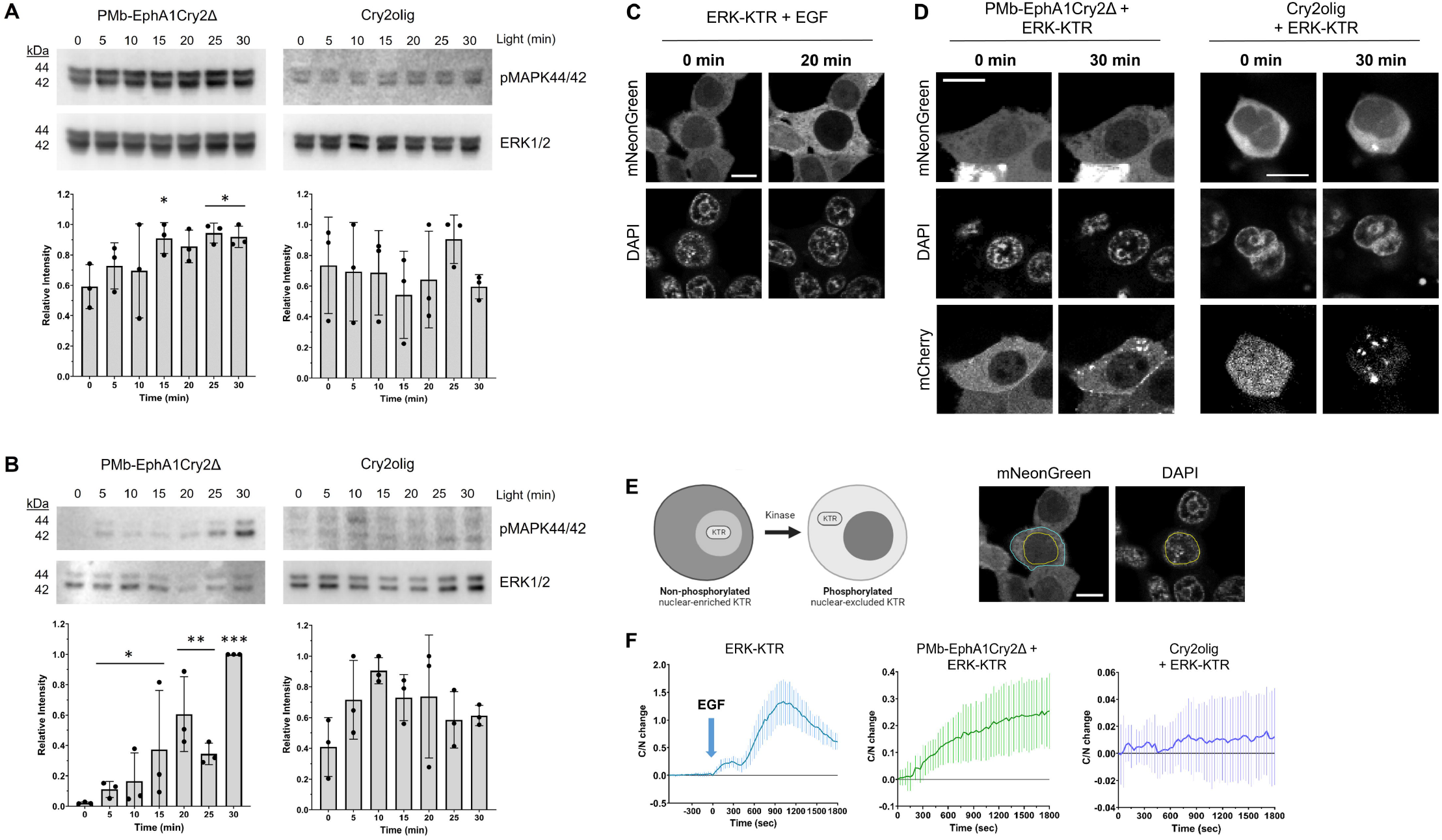
ERK phosphorylation in response to EphA1Cry2Δ activation. **(A,B)** Western blots of PMb-EphA1Cry2Δ (left) and Cry2olig (right) transfected HEK293T cell lysates over 5 min time intervals of light stimulation, immunostained for ERK phosphorylation (pMAPK44/42) and ERK1/2 loading control. **(A)** Cells were not serum-starved prior to lysis. **(B)** Cells were serum-starved for 2 hours prior to light exposure and lysis. **(A**,**B)** Bottom graphs: Quantitation of normalized pMAPK44/42 band intensity to ERK1/2. Error bars represent standard deviation, n=3 (unpaired t-test to 0 min time point; ^*^p < 0.05, ^**^p < 0.01, ^***^p < 0.001). **(C, D)** Confocal images of HEK293T cells transfected with **(C)** only ERK-KTR with 500 ng/mL EGF stimulus, **(D)** ERK-KTR + PMb-EphA1Cry2Δ, and ERK-KTR + Cry2olig taken pre (0 min) and post blue light (480 nm, 30 sec intervals, over 30 min). Images collected on DAPI channel (nuclear stain), mNeonGreen channel (ERK-KTR), and for (D) mCherry channel (PMb-EphA1Cry2Δ and Cry2olig). Scale bar: 10 µm. **(E)** ERK-KTR is localized to the nucleus in its non-phosphorylated state. Endogenous ERK activation upon kinase activity leads to nuclear translocation, which in turn translocates ERK-KTR out of the nucleus into the cytosol. Right images: Example quantification using Fiji ImageJ to measure the fluorescent intensity of the cytosol (cyan ROI) and nucleus (yellow ROI). **(F)** Quantitation of fluorescent intensity of the cytosol divided by the intensity within the nucleus (C/N) change over time. ERK-KTR (B) included 10 min pre-stimulus imaging. Error bars represent standard deviation; n=6 cells.

### EphA1Cry2 activation in differentiated Neuro2a cells

We also investigated the effects of PMb-EphA1Cry2Δ activation in other cell types other than HEK293T cells. Neuro2a (N2a) cells were selected as the N2a cell line is a mouse neural crest-derived cell line commonly used to study neuronal differentiation and its related signaling pathways (Tremblay et al., 2010). Differentiated N2a cells transfected with PMb-EphA1Cry2Δ show reversible, localized clustering at the membrane of the neuronal-like processes (**Fig. 5A**). Small protrusions along these processes were identified at time 0 min of the PMb-EphA1Cry2Δ transfected cells and retract into the processes 10 min after light exposure. These protrusions returned after 10 min in the dark condition (**Fig. 5B; Supporting Movie 3**). Based on these results, the EphA1Cry2 construct could be applied for the induction of morphological changes relevant to synaptic structural plasticity.

**Figure 5.**
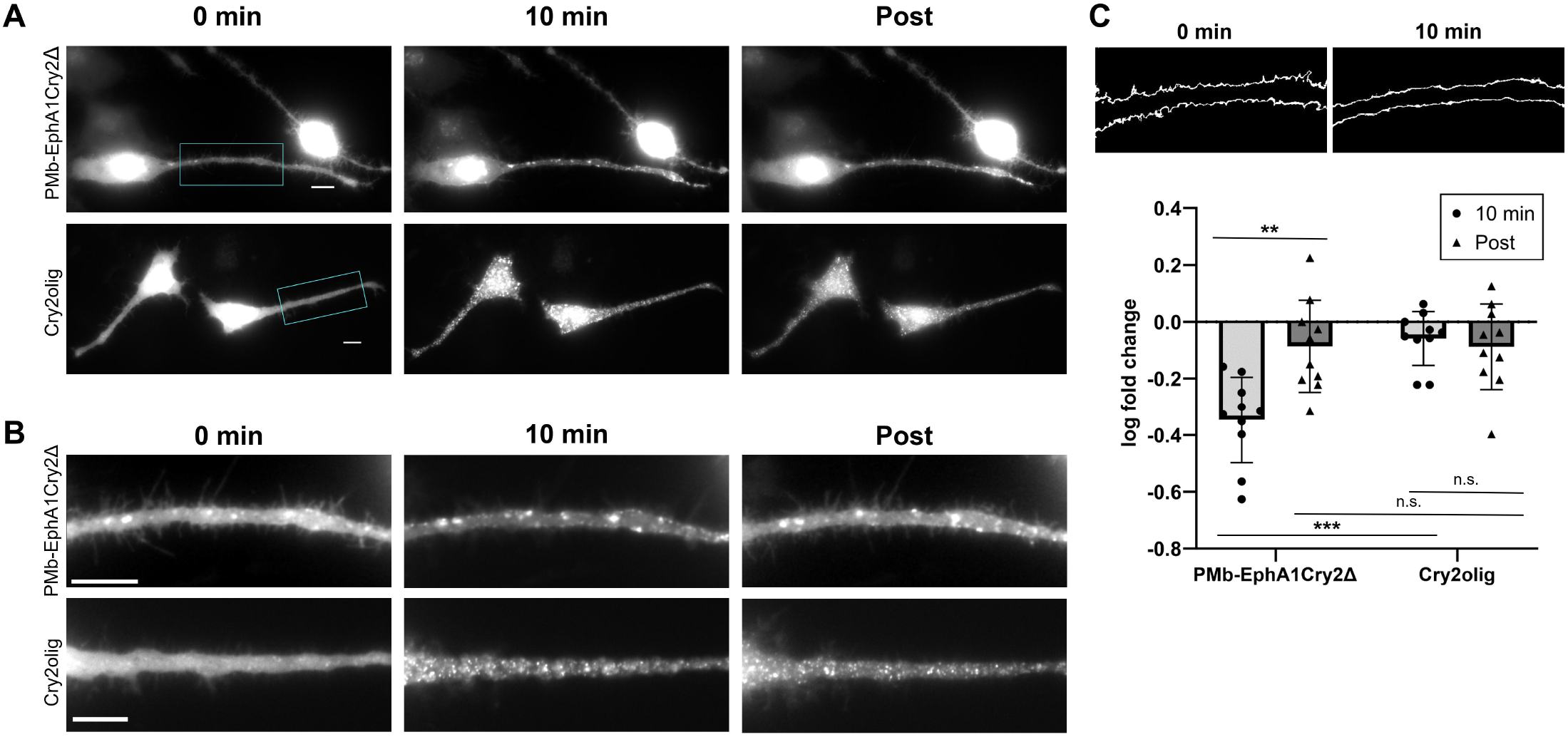
Optogenetic clustering of transfected N2a cells. (A) Widefield images of differentiated N2a cells transfected with PMb-EphA1Cry2Δ and Cry2olig taken pre-light exposure (0 min), after 10 min of blue light exposure (480 nm, 50 ms exposure, 30 sec intervals), and 10 min post-light exposure (Post). Images collected on mCherry channel (200 ms exposure, 553 nm). Scale bar: 10 µm. **(B)** Zoomed region of interest (cyan box) of neuronal-like process of a cell in (A). Scale bar: 10 µm. **(C)** Top: Threshold images of PMb-EphA1Cry2Δ transfected cells (at 0 min and 10 min of light exposure) from (B) used to count protrusions along neuronal-like processes of differentiated N2a cells. Bottom: Log fold change in protrusion number on the neuronal-like processes. 10 min of light exposure and post-light exposure were quantified in respect to protrusion number at time 0 min of light exposure. Error bars represent standard deviation, n=10 cells (unpaired t-test; ^***^p < 0.0001,^**^p < 0.01, n.s. = not significant).

### Summary and Conclusions

Eph receptors are the key to many signaling pathways that regulate cell-cycle progression and cell migration (Liang et al., 2019). In our studies, we designed constructs of an optogenetic EphA1 receptor using Cry2olig that clustered in response to blue light stimulus. We found that this light-initiated clustering resulted in activation by auto-phosphorylation of the EphA1 receptor’s tyrosine kinase domain and induced downstream ERK signaling in HEK293T cells. We also found that expression of our PMb-EphA1Cry2Δ construct in N2a cells resulted in modulation of protrusive activity along the neuronal-like processes of the cells.

Previous studies on EphA1 receptor have elucidated its role in cell mobility and morphology (Ieguchi & Maru, 2019; Yamazaki et al., 2009). Specifically, ephrinA1 stimulation of EphA1 receptor resulted in activation of the RhoA/ROCK pathway, which promotes cell rounding and migration (Yamazaki et al., 2009). Rho-GTPases are associated with the integrin-linked kinase (ILK) pathways and help regulate the actin cytoskeleton through GTP binding and hydrolysis (Yamazaki et al., 2009). Of the Rho GTPases, RhoA regulates actomyosin contractility and stress-fiber formation, and Rac1 regulates lamellipodia formation by the WAVE-mediated ARP2/3 complex (Beco et al., 2018; Kunschmann et al., 2019). EphA1 has been shown to interact with ILK, a kinase that binds to integrin and promotes signaling transduction from the ECM to the cytosol. Activated EphA1 inhibits ILK activity, which inactivates Rac1, thereby inhibiting cell spreading in an ephrin-dependent manner (Yamazaki et al., 2009). Removal of the SAM domain prevented the binding of ILK and eliminated the cell spreading defect (Yamazaki et al., 2009). Conversely, studies of the SAM domain in EphA2 receptor found that a deletion of the SAM domain results in an increase in the ability of the receptor to dimerize and activate even in the absence of ephrinA1-Fc stimulation (Singh et al., 2017). A ΔSAM mutant of EphA1 receptor specifically can still be activated through its kinase domain but associated morphological change were inhibited (Yamazaki et al., 2009). Interestingly, our results indicate that activation of receptor tyrosine kinase domains via EphA1Cry2olig clustering can induce cell rounding with and without the presence of the SAM domain. While a pathway of EphA1 signaling has been proposed (Ieguchi & Maru, 2019), further elucidation of EphA1 signaling pathways is still needed to determine the effect of EphA1 activity. The benefit of our optogenetic constructs is that the EphA1 signaling is simplified to include only forward signaling, as the extracellular regions have been removed.

Other studies in EphA receptor signaling reported ephrinA1-induced EphA receptor activation resulted in an inhibition of the Ras/Raf/MEK/MAPK pathway that regulates cell proliferation (Miao et al., 2000, 2001). These studies were also conducted in HEK293T cells; however, EphA2 receptor is the predominate endogenously expressed EphA receptor in HEK293T cells and the other cell types used (Miao et al., 2001). Our light-activated PMb-EphA1Cry2Δ construct induced ERK phosphorylation, which upregulates proliferation (Mebratu & Tesfaigzi, 2009). This finding may not be surprising as EphA1 receptor has been found to up-regulated in certain types of cancer (Ieguchi & Maru, 2019). As EphA1 receptors are essential for angiogenesis, tumors may increase expression of EphA1 receptor to promote proliferation. Coupled with a decrease in cell adhesion and an increase in cell migration, EphA1 receptor-activated proliferation would induce metastasis of a tumor cell if EphA1 was overexpressed (Ieguchi & Maru, 2019). EphA1 expression is up-regulated by the activating transcription factor 3 (ATF3), which has been found to be induced under stress conditions (Masuda et al., 2008). A study of expression patterns in clear cell renal cell carcinoma (ccRCC) consistently showed that a lower EphA1 expression in tumor cells corresponded to a more positive prognosis (Toma et al., 2014). In hepatocellular carcinoma (HCC), a common liver tumor, up-regulated EphA1 promotes Akt/mTOR-activated transcription of the chemokine, stromal cell-derived factor 1 (SDF-1). SDF-1 is then secreted by the tumor cell and binds to the CXCR4 receptor on an endothelial progenitor cell, leading to chemotaxis and tumorigenesis (Y. Wang et al., 2016).

Although there have been few studies on EphA receptors in neurons, the research on their ligand, ephrinA1, has suggested that stimulation of ephrinA1 results in neuron migration (Bravo-Cordero et al., 2013). One of the major EphA receptors in neurons is EphA4, a homolog of EphA1 and EphA2, that has been shown to interact with EphA1 (Murai et al., 2003). Through binding of ephrinA3 on a neighboring astrocyte, EphA4 forward signaling on a dendritic spine can decrease its spine length and width (Murai et al., 2003). This may also be the case with PMb-EphA1Cry2Δ, as the construct’s activation resulted in a decrease of protrusions along the processes of N2a cells. In primary neurons, EphA1 activation could result in a decrease in spines which would prime the neuron for deadhesion and migration (Holen et al., 2010; Murai et al., 2003). As Eph receptors are integral in cell-cell communication during neurogenesis, their dysfunction can play a role in neurological disorders and neurodegenerative diseases (Liang et al., 2019). Recently, researchers have found that Alzheimer’s disease (AD) pathology could be detected via biomarkers in a patient’s genome (H. F. Wang et al., 2015). In an analysis of recent genome studies, the EphA1 locus was found to be highly associated with AD (Carrasquillo et al., 2011; Naj et al., 2011; H. F. Wang et al., 2015). The A-allele of EphA1 was correlated with a decrease in hippocampal atrophy in mild cognitive impairment patients and was associated with a higher glucose cerebral metabolic rate in certain parts of the brain (H. F. Wang et al., 2015). Understanding the role EphA1 plays in different cell types, including neurons, could yield new insights into cancer and neurodegenerative diseases.

## Author contributions

A. I. W., K. Z., and R. M. H. methodology; A. I. W. formal analysis; A. I. W. and K. Z. investigation; A. I. W. and R. M. H. writing.

## Funding and additional information

This work was supported by the National Institutes of Health (grant no.: 1R15NS125564-01). The content is solely the responsibility of the authors and does not necessarily represent the official views of the National Institutes of Health.

## Supporting Figures

**Supporting Figure 1. Spatiotemporal light activation of EphA1Cry2 constructs. (A)** Confocal images of HEK293T cells transfected with Cry2olig, PMa-EphA1Cry2, PMa-EphA1Cry2Δ, or PMb-EphA1Cry2Δ taken pre-illumination (0 min) and after blue light exposure (480 nm, 30 sec intervals for 10 min) only inside circular region (yellow). Images collected on mCherry channel (200 ms exposure, 553 nm). Scale bar: 10 µm. Right column: Quantitation of the relative intensity of inside and outside the region of interest (ROI) over blue light exposure starting at 90 sec (blue arrow). Error bars represent standard deviation; n=3 regions (unpaired t-test,^*^p < 0.05). **(B)** Widefield images of HEK293T cells transfected with Cry2olig, PMa-EphA1Cry2Δ, or PMb-EphA1Cry2Δ before light exposure (0 min), after 5 min of light exposure (480 nm, 50 ms exposure, 30 sec intervals), and 10 min post-light exposure (Post). Images collected on mCherry channel (200 ms exposure, 553 nm). Scale bar: 10 µm.

**Supporting Figure 2. Cell rounding assay in response to EphA1Cry2 light activation. (A)** Widefield images of HEK293T cells transfected with Cry2olig, PMa-EphA1Cry2, PMa-EphA1Cry2Δ, or PMb-EphA1Cry2Δ before light exposure (0 min) and after 5 min of light exposure (480 nm, 50 ms exposure, 30 sec intervals). Images collected on mCherry channel (553 nm, 200 ms exposure). Cell areas were measured by tracing the cell around the membrane (yellow) using Fiji ImageJ. Middle column shows the trace of the cell area (yellow) at time 0 min overlaid with the cell at time 5 min. Scale bar: 10 µm. **(B)** Quantitation of the percent area change (% change in the cell area at 10 min over cell area at 0 min) of transfect HEK293T cells in (A). Error bars represent standard deviation, n=6 cells (unpaired t-test; ^***^p < 0.001).

## Supporting Movie

**Supporting Movie 1. Fluorescence widefield microscopy of PMb-EphA1Cry2**Δ **light activation in HEK293T cells**. Whole cell activation of PMb-EphA1Cry2Δ results in construct clustering at the membrane in response to blue light, which is reversible after removal of blue light. Images acquired every 30 s (553 nm; 200 ms exposure). Blue light activation (480 nm, 50 ms pulse) protocol: 0:00 to 5:30 min:s (blue light pulse every 30 s); 6:00 to 16:00 min:s (no blue light). Scale bar: 10 µm.

**Supporting Movie 2. Confocal microscopy of PMb-EphA1Cry2**Δ **light activation with ERK-KTR and DAPI nuclear stain in HEK293T cells**. Simultaneous PMb-EphA1Cry2Δ activation and imaging of ERK-KTR with 488 nm laser (5% power) occurs every 30 s over 30 min. Left panel: mCherry channel showing PMb-EphA1Cry2Δ activation. Middle panel: DAPI channel showing stain of nucleus. Right panel: mNeonGreen channel showing ERK-KTR translocation. Images acquired every 30 s (553 nm; 5% power). Scale bar: 10 µm.

**Supporting Movie 3. Fluorescence widefield microscopy of PMb-EphA1Cry2**Δ **light activation in the neuronal-like processes of Neuro2A cells**. Images acquired every 30 s (553 nm; 200 ms exposure). Blue light activation (480 nm, 50 ms pulse) protocol: 0:00 to 5:00 min:s (no blue light); 5:30 – 15:30 min:s (blue light pulse every 30 s); 16:00 – end min:s (no blue light). Scale bar: 10 µm.

## Notes

### Competing Interest Statement

The authors have declared no competing interest.

### Summary of Updates

Updates to Figure 4 (serum starved Erk western blot added); minor changes to text.

## References

Arvanitis, D., & Davy, A. (2008). Eph/ephrin signaling: Networks. In Genes and Development (Vol. 22, Issue 4). 10.1101/gad.1630408

Barquilla, A., & Pasquale, E. B. (2015). Eph receptors and ephrins: Therapeutic opportunities. In Annual Review of Pharmacology and Toxicology (Vol. 55). 10.1146/annurev-pharmtox-011112-140226

Beco, S. de, Vaidžiulytė, K., Manzi, J., Dalier, F., Federico, F. di, Cornilleau, G., Dahan, M., & Coppey, M. (2018). Optogenetic dissection of Rac1 and Cdc42 gradient shaping. Nature Communications, 9(1). 10.1038/s41467-018-07286-8

Bravo-Cordero, J. J., Magalhaes, M. A. O., Eddy, R. J., Hodgson, L., & Condeelis, J. (2013). Functions of cofilin in cell locomotion and invasion. In Nature Reviews Molecular Cell Biology (Vol. 14, Issue 7). 10.1038/nrm3609

Bugaj, L. J., Spelke, D. P., Mesuda, C. K., Varedi, M., Kane, R. S., & Schaffer, D. V. (2015). Regulation of endogenous transmembrane receptors through optogenetic Cry2 clustering. Nature Communications, 6. 10.1038/ncomms7898

Carrasquillo, M. M., Belbin, O., Hunter, T. A., Ma, L., Bisceglio, G. D., Zou, F., Crook, J. E., Pankratz, V., Sando, S. B., Aasly, J. O., Barcikowska, M., Wszolek, Z. K., Dickson, D. W., Graff-Radford, N. R., Petersen, R. C., Passmore, P., Morgan, K., & Younkin, S. G. (2011). Replication of EPHA1 and CD33 associations with late-onset Alzheimer’s disease: A multi-centre case-control study. Molecular Neurodegeneration, 6(1). 10.1186/1750-1326-6-54

Chavez-Abiega, S., Gönloh, M. L. B., Gadella, T. W. J., Bruggeman, F. J., & Goedhart, J. (2022). Single-cell imaging of ERK and Akt activation dynamics and heterogeneity induced by G-protein-coupled receptors. Journal of Cell Science, 135(6). 10.1242/jcs.259685

Darling, T. K., & Lamb, T. J. (2019). Emerging roles for Eph receptors and ephrin ligands in immunity. In Frontiers in Immunology (Vol. 10, Issue JULY). 10.3389/fimmu.2019.01473

Dessauges, C., Mikelson, J., Dobrzyński, M., Jacques, M., Frismantiene, A., Gagliardi, P. A., Khammash, M., & Pertz, O. (2022). Optogenetic actuator – ERK biosensor circuits identify MAPK network nodes that shape ERK dynamics. Molecular Systems Biology, 18(6), e10670. 10.15252/msb.202110670

Duan, L., Hope, J., Ong, Q., Lou, H. Y., Kim, N., McCarthy, C., Acero, V., Lin, M. Z., & Cui, B. (2017). Understanding CRY2 interactions for optical control of intracellular signaling. Nature Communications, 8(1). 10.1038/s41467-017-00648-8

Gucciardo, E., Sugiyama, N., & Lehti, K. (2014). Eph- and ephrin-dependent mechanisms in tumor and stem cell dynamics. In Cellular and molecular life sciences: CMLS (Vol. 71, Issue 19). 10.1007/s00018-014-1633-0

Hernández-Candia, C. N., Pearce, S., & Tucker, C. L. (2021). A modular tool to query and inducibly disrupt biomolecular condensates. Nature Communications, 12(1). 10.1038/s41467-021-22096-1

Hirai, H., Maru, Y., Hagiwara, K., Nishida, J., & Takaku, F. (1987). A novel putative tyrosine kinase receptor encoded by the eph gene. Science, 238(4834). 10.1126/science.2825356

Holen, H. L., Nustad, K., & Aasheim, H. C. (2010). Activation of EphA receptors on CD4+CD45RO+ memory cells stimulates migration. Journal of Leukocyte Biology, 87(6). 10.1189/jlb.0709497

Hughes, R. M. (2018). A compendium of chemical and genetic approaches to light-regulated gene transcription. In Critical Reviews in Biochemistry and Molecular Biology (Vol. 53, Issue 5). 10.1080/10409238.2018.1487382

Hughes, R. M., & Virag, J. A. I. (2020). Harnessing the power of eph/ephrin biosemiotics for theranostic applications. In Pharmaceuticals (Vol. 13, Issue 6). 10.3390/ph13060112

Ieguchi, K., & Maru, Y. (2019). Roles of EphA1/A2 and ephrin-A1 in cancer. In Cancer Science (Vol. 110, Issue 3). 10.1111/cas.13942

Kennedy, M. J., Hughes, R. M., Peteya, L. A., Schwartz, J. W., Ehlers, M. D., & Tucker, C. L. (2010). Rapid blue-light-mediated induction of protein interactions in living cells. Nature Methods, 7(12). 10.1038/nmeth.1524

Kunschmann, T., Puder, S., Fischer, T., Steffen, A., Rottner, K., & Mierke, C. T. (2019). The Small GTPase Rac1 Increases Cell Surface Stiffness and Enhances 3D Migration Into Extracellular Matrices. Scientific Reports, 9(1). 10.1038/s41598-019-43975-0

Lai, K.-O., & Ip, N. Y. (2009). Synapse development and plasticity: Roles of ephrin/Eph receptor signaling. Current Opinion in Neurobiology, 19(3), 275–283. 10.1016/j.conb.2009.04.009

Liang, L. Y., Patel, O., Janes, P. W., Murphy, J. M., & Lucet, I. S. (2019). Eph receptor signalling: From catalytic to non-catalytic functions. In Oncogene (Vol. 38, Issue 39). 10.1038/s41388-019-0931-2

Lisabeth, E. M., Falivelli, G., & Pasquale, E. B. (2013). Eph receptor signaling and ephrins. Cold Spring Harbor Perspectives in Biology, 5(9). 10.1101/cshperspect.a009159

Locke, C., Machida, K., Wu, Y., & Yu, J. (2017). Optogenetic activation of EphB2 receptor in dendrites induced actin polymerization by activating Arg kinase. Biology Open, 6(12). 10.1242/bio.029900

Masuda, J., Usui, R., & Maru, Y. (2008). Fibronectin type I repeat is a nonactivating ligand for EphA1 and inhibits ATF3-dependent angiogenesis. The Journal of Biological Chemistry, 283(19), 13148–13155. 10.1074/jbc.M702164200

Mebratu, Y., & Tesfaigzi, Y. (2009). How ERK1/2 activation controls cell proliferation and cell death: Is subcellular localization the answer? Cell Cycle (Georgetown, Tex.), 8(8), 1168–1175. 10.4161/cc.8.8.8147

Miao, H., Burnett, E., Kinch, M., Simon, E., & Wang, B. (2000). Activation of EphA2 kinase suppresses integrin function and causes focal-adhesion-kinase dephosphorylation. Nature Cell Biology, 2(2), 62–69. 10.1038/35000008

Miao, H., & Wang, B. (2012). EphA receptor signaling—Complexity and emerging themes. Seminars in Cell & Developmental Biology, 23(1), 16–25. 10.1016/j.semcdb.2011.10.013

Miao, H., Wei, B.-R., Peehl, D. M., Li, Q., Alexandrou, T., Schelling, J. R., Rhim, J. S., Sedor, J. R., Burnett, E., & Wang, B. (2001). Activation of EphA receptor tyrosine kinase inhibits the Ras/MAPK pathway. Nature Cell Biology, 3(5), 527–530. 10.1038/35074604

Murai, K. K., Nguyen, L. N., Irie, F., Yu, Y., & Pasquale, E. B. (2003). Control of hippocampal dendritic spine morphology through ephrin-A3/EphA4 signaling. Nature Neuroscience, 6(2). 10.1038/nn994

Naj, A. C., Jun, G., Beecham, G. W., Wang, L. S., Vardarajan, B. N., Buros, J., Gallins, P. J., Buxbaum, J. D., Jarvik, G. P., Crane, P. K., Larson, E. B., Bird, T. D., Boeve, B. F., Graff-Radford, N. R., Jager, P. L. D., Evans, D., Schneider, J. A., Carrasquillo, M. M., Ertekin-Taner, N., … Schellenberg, G. D. (2011). Common variants at MS4A4/MS4A6E, CD2AP, CD33 and EPHA1 are associated with late-onset Alzheimer’s disease. Nature Genetics, 43(5). 10.1038/ng.801

Resh, M. D. (1999). Fatty acylation of proteins: New insights into membrane targeting of myristoylated and palmitoylated proteins. Biochimica Et Biophysica Acta, 1451(1), 1–16. 10.1016/s0167-4889(99)00075-0

Singh, D. R., Ahmed, F., Paul, M. D., Gedam, M., Pasquale, E. B., & Hristova, K. (2017). The SAM domain inhibits EphA2 interactions in the plasma membrane. Biochimica et Biophysica Acta - Molecular Cell Research, 1864(1). 10.1016/j.bbamcr.2016.10.011

Taslimi, A., Vrana, J. D., Chen, D., Borinskaya, S., Mayer, B. J., Kennedy, M. J., & Tucker, C. L. (2014). An optimized optogenetic clustering tool for probing protein interaction and function. Nature Communications, 5. 10.1038/ncomms5925

Thaa, B., Biasiotto, R., Eng, K., Neuvonen, M., Götte, B., Rheinemann, L., Mutso, M., Utt, A., Varghese, F., Balistreri, G., Merits, A., Ahola, T., & McInerney, G. M. (2015). Differential Phosphatidylinositol-3-Kinase-Akt-mTOR Activation by Semliki Forest and Chikungunya Viruses Is Dependent on nsP3 and Connected to Replication Complex Internalization. Journal of Virology, 89(22), 11420–11437. 10.1128/JVI.01579-15

Toma, M. I., Erdmann, K., Diezel, M., Meinhardt, M., Zastrow, S., Fuessel, S., Wirth, M. P., & Baretton, G. B. (2014). Lack of ephrin receptor A1 is a favorable independent prognostic factor in clear cell renal cell carcinoma. PLoS ONE, 9(7). 10.1371/journal.pone.0102262

Tremblay, R. G., Sikorska, M., Sandhu, J. K., Lanthier, P., Ribecco-Lutkiewicz, M., & Bani-Yaghoub, M. (2010). Differentiation of mouse Neuro 2A cells into dopamine neurons. Journal of Neuroscience Methods, 186(1). 10.1016/j.jneumeth.2009.11.004

Wang, H. F., Tan, L., Hao, X. K., Jiang, T., Tan, M. S., Liu, Y., Zhang, D. Q., & Yu, J. T. (2015). Effect of EPHA1 genetic variation on cerebrospinal fluid and neuroimaging biomarkers in healthy, mild cognitive impairment and Alzheimer’s disease cohorts. Journal of Alzheimer’s Disease, 44(1). 10.3233/JAD-141488

Wang, Y., Yu, H., Shan, Y., Tao, C., Wu, F., Yu, Z., Guo, P., Huang, J., Li, J., Zhu, Q., Yu, F., Song, Q., Shi, H., Zhou, M., & Chen, G. (2016). EphA1 activation promotes the homing of endothelial progenitor cells to hepatocellular carcinoma for tumor neovascularization through the SDF-1/CXCR4 signaling pathway. Journal of Experimental and Clinical Cancer Research, 35(1). 10.1186/s13046-016-0339-6

Yamazaki, T., Masuda, J., Omori, T., Usui, R., Akiyama, H., & Maru, Y. (2009). EphA1 interacts with integrin-linked kinase and regulates cell morphology and motility. Journal of Cell Science, 122(2). 10.1242/jcs.036467

Yang, J. S., Wei, H. X., Chen, P. P., & Wu, G. (2018). Roles of Eph/ephrin bidirectional signaling in central nervous system injury and recovery (Review). In Experimental and Therapeutic Medicine (Vol. 15, Issue 3). 10.3892/etm.2018.5702

