## Supplementary figures and images for "Optogenetic Regulation of EphA1 RTK Activation and Signaling"

### Supporting Figures 1 and 2

**SUPPORTING FIGURE 1**

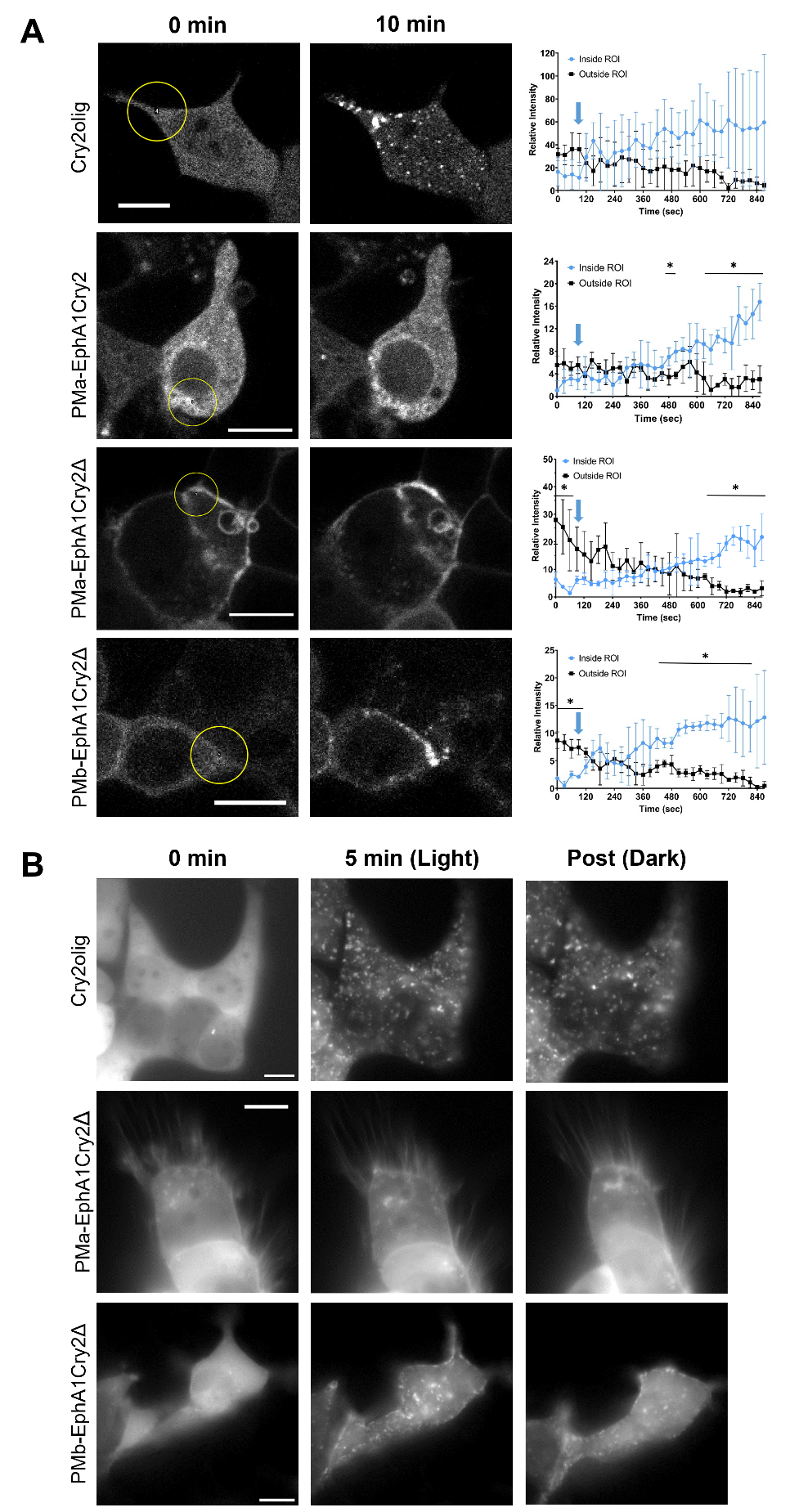

**SUPPORTING FIGURE 2**
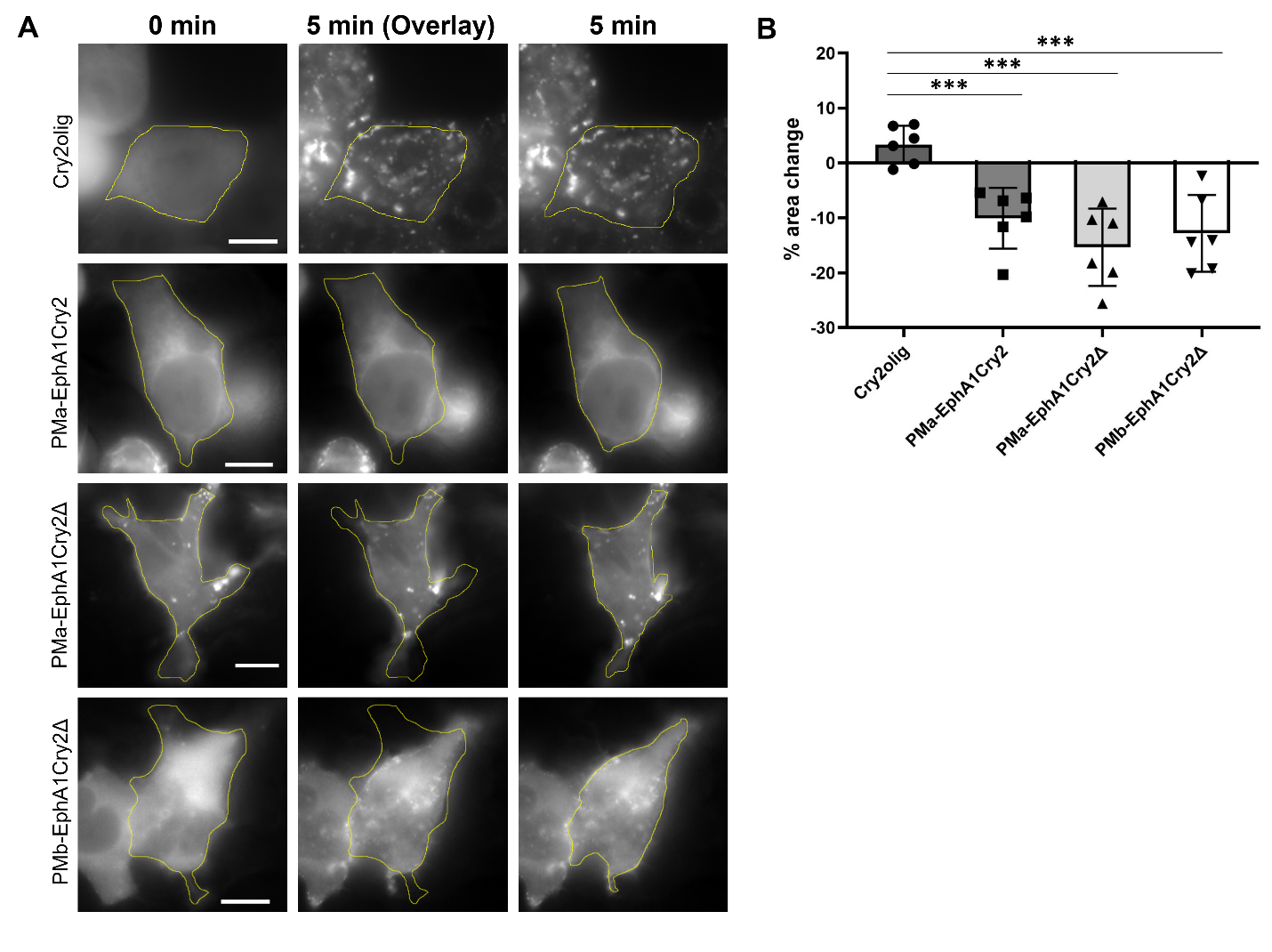
